# Genome Stability under Silence: DNA Repair Networks in Quiescent Fission Yeast

**DOI:** 10.1101/2025.08.24.671962

**Authors:** Samia Miled

## Abstract

Most of our current understanding of genome integrity derives from studies in proliferating cells, yet most somatic cells in multicellular organisms reside in non-dividing, quiescent states. Using *Schizosaccharomyces pombe*, we dissected the mechanisms by which quiescent cells maintain genome stability in the absence of DNA replication. Combining time-resolved mutational analyses, DNA damage assays, and genetic dissection of DNA repair pathways, we found that quiescent cells progressively accumulate distinct types of spontaneous lesions-particularly uracil residues, abasic sites, and ribonucleotide insertions-which are counteracted by a modular network of repair mechanisms.

Base excision repair (BER), ribonucleotide excision repair (RER), and R-loop resolution each contribute uniquely to genome surveillance in G0. We show that uracil incorporation becomes a predominant threat under quiescent conditions, especially when nucleotide pools are imbalanced. BER-deficient mutants (e.g., *nth1Δ*, *ung1Δ*) exhibit mutation spectra dominated by C: G > T: A transitions and oxidative lesions, while synthetic combinations reveal compensatory or epistatic interactions. Using single-cell micromanipulation and viability assays, we show that specific gene deletions (e.g., *hnt3Δrhp52Δ*, *sen1Δrad13Δ*) severely compromise post-quiescence recovery, underscoring the importance of cooperative DNA repair even in non-replicative contexts.

Our results delineate a functionally compartmentalized hierarchy of DNA repair activities during quiescence, providing a new framework to understand how non-dividing cells limit genome instability, with implications for aging, cancer dormancy, and neurodegeneration.

## INTRODUCTION

Eukaryotic cellular quiescence, a non-dividing, reversible G₀ state, is a strategy to conserve energy and preserve tissue homeostasis while retaining the capacity for reactivation. In multicellular organisms, the majority of differentiated and adult cells - such as neurons, hepatocytes, hematopoietic stem cells, and immune cells - persist predominantly in this quiescent state [1,2].

Contrary to the classical notion that quiescent cells are metabolically inert and genetically stable, growing evidence indicates they endure continuous endogenous stress. Residual transcriptional activity, mitochondrial reactive oxygen species, and metabolic fluctuations impose persistent damage on genomic DNA [3,4]. Since quiescent cells lack DNA replication, they cannot rely on replication-coupled repair and instead depend on replication-independent pathways to safeguard genome integrity.

*Schizosaccharomyces pombe* (fission yeast) serves as a highly informative model for dissecting these mechanisms. Quiescence can be systematically induced by nitrogen deprivation, leading to ordered G₀ arrest, and the small, well-annotated genome enables fine-scaled mutational and genetic studies [5,6]. Quiescent *S. pombe* cells remain viable for extended periods, remain metabolically active, and maintain DNA damage responses, yet they progressively accumulate spontaneous mutations, establishing a quiescence-specific mutational signature [7].

Despite this, the lesion types driving these mutations and their repair dependencies are not well defined. Emerging evidence points to multiple repair pathways - such as base excision repair (BER), ribonucleotide excision repair (RER), single-strand break processing, nucleotide excision repair (NER), and R-loop resolution - as active contributors to genome maintenance in G0. For instance, BER and NER collaborate synergistically to support survival in stationary-phase fission yeast [59]. Additionally, Ku/NHEJ and MRN complex proteins are essential for handling DNA breaks during G0, while NER also contributes to repair of spontaneous lesions over time [53].

Our study aims to delineate how quiescent genomes in *S. pombe* are maintained by coordinated and lesion-specific repair mechanisms. We first examine viability and mutation accrual in wild-type cells during extended G₀. We then employ genetic deletions to dissect the roles of repair pathways - such as BER (*Ung1, Nth1*), RER (*Rnh201, Hnt3*), topoisomerase-linked end processing (*Tdp1*), and RNA–DNA hybrid remediation (*Sen1*). By integrating molecular lesion detection (e.g., uracil quantification via LC–MS/MS), mutational spectrum analysis, and single-cell G₀ exit profiling, we uncover a modular repair network that preserves genome fidelity in the absence of replication.

## Materials & Methods

### Strains and culture conditions

*Schizosaccharomyces pombe* strains used in this study are derivatives of the wild-type 972 h-background. Gene deletions were performed using PCR-based homologous recombination with kanMX6, resistance cassettes. Mutants were verified by PCR and phenotypic assays.

Quiescence (G0) was induced by nitrogen starvation as described (Marguerat et al., 2012). Briefly, logarithmically growing cultures in YES were filtered and transferred into EMM minus nitrogen (EMM–N) at OD600 = 0.5. Viability was monitored by colony-forming unit (CFU) counting after plating serial dilutions on YES.

### Strain construction and annotation

Deletion mutants were generated by PCR-mediated gene replacement using a kanMX6 cassette and standard homologous recombination techniques. All double and triple mutants were confirmed by colony PCR and viability assays.

### FOA mutagenesis assay

Mutation frequency was measured by selection for resistance to 5-fluoroorotic acid (5-FOA; Thermo Fisher). At each indicated time point in quiescence (Day 1 to Day 30), 1 × 10⁷ cells were plated onto YES + FOA (1 mg/mL) and YES control plates. Colony counts were performed after 3–5 days at 30 °C. Mutation rate per day was calculated as the slope of FOA-resistant colonies accumulation over time.

Complementation assays using integrative *ura4+* and *ura5+* plasmids allowed assignment of resistance to specific gene inactivation. Mutational spectra were determined by amplifying *ura4* and *ura5* loci by PCR, gel-purification, and Sanger sequencing (Eurofins Genomics). Sequence alignments were performed using SnapGene.

### Mutation Screening and Sequence Analysis

Spontaneous loss-of-function mutations in *ura4* and *ura5* were identified by selection of FOA-resistant (FOA^R^) clones during quiescence. Cells were plated on YES medium supplemented with 1 mg/mL 5-fluoroorotic acid (5-FOA). FOA^R^ colonies were picked after 3–5 days of incubation at 30 °C.

To distinguish whether mutations affected *ura4* or *ura5*, multiple independent FOA^R^ colonies were re-streaked on rich YP plates and incubated at 30 °C. Mutants with *ura5* loss-of-function appear as red colonies due to accumulation of a red pigment, while *ura4* mutants retain a white or cream phenotype. This phenotypic differentiation allowed rapid assignment of the mutated gene in most cases.

Genomic DNA was extracted from individual FOA^R^ clones using a standard phenol–chloroform protocol. The corresponding *ura4* or *ura5* loci were PCR-amplified using gene-specific forward and reverse primers. PCR products were visualized by agarose gel electrophoresis, purified, and subjected to Sanger sequencing.

Mutations were identified by alignment to the *Schizosaccharomyces pombe* reference genome (PomBase identifiers: *ura4*– SPAC1093.11c; *ura5* – SPAC3F10.07c), and categorized as single-nucleotide substitutions, insertions, deletions, duplications, or complex structural events, using SNAPGENE software version 8.1.1.

### DNA damage detection assays

To detect uracil-derived lesions, total DNA was extracted from cells at Days 2, 4, and 8 in G0 using the MasterPure DNA purification kit (Lucigen). Aliquots of 1 μg DNA were incubated with recombinant Ung1 and EndoVIII (New England Biolabs) for 30 min at 37 °C, and analyzed by agarose gel electrophoresis. Fragmentation indicates unrepaired uracil and oxidized base damage.

ARP–dot blot was used to detect AP sites following uracil excision. Membrane-bound DNA was incubated with Aldehyde Reactive Probe (ARP; Dojindo), followed by streptavidin–HRP and chemiluminescence detection using luminol substrate. Signal intensity was quantified with ImageJ.

### Drop test and microcolony growth assay

5 μL of 5-fold serial dilutions of G0 cultures were spotted onto YES and YES+FOA plates. Plates were incubated at 30 °C and photographed after 3–5 days. Microcolony formation was scored under the dissecting microscope.

### Statistical analysis

All experiments were performed with at least three independent biological replicates. Data are presented as mean ± SEM. Statistical analyses were performed using GraphPad Prism 10 or R. Significance was assessed by Student’s t-test or ANOVA as appropriate. Mutation spectra differences were analyzed using Fisher’s exact test or chi-square tests.

### Uracil Detection by LC/MS and Mass Spectrometry

Genomic DNA was extracted from quiescent *Schizosaccharomyces pombe* cells aged in G0 for 8 days, using a phenol-chloroform purification protocol. Quantification of free uracil residues was performed by liquid chromatography coupled to mass spectrometry (LC/MS).

Samples were analyzed on an Agilent 1290 Infinity II UHPLC system coupled to an Agilent 6530 Q-TOF mass spectrometer, operating in positive ionization mode. Chromatographic separation was carried out using a reverse-phase C18 column (2.1 × 50 mm, 1.8 μm particle size) at 35 °C. The mobile phase consisted of water (0.1% formic acid) and acetonitrile (0.1% formic acid) in a gradient elution profile. Injection volume was 5 μL. Mass spectra were acquired in full-scan MS mode, and uracil was identified based on accurate mass (m/z 113.03 [M+H]+), retention time, and comparison with authentic uracil standards. Data were processed using Agilent software, and relative quantification was based on peak area integration.

## Results

### 1. Quiescent fission yeast cells progressively lose viability and accumulate base lesions in a pathway-dependent manner

In multicellular organisms, the majority of somatic cells reside in a non-dividing state known as quiescence (G0), a reversible arrest phase required for tissue homeostasis, regeneration, and longevity [1,2]. Despite the absence of DNA replication, quiescent cells remain metabolically active and are continuously exposed to endogenous genotoxic stress, including oxidative damage, spontaneous base deamination, and transcription-associated DNA structures [29,30,20]. Yet, the dynamics and origin of genome instability during G0 remain poorly understood, in part due to the difficulty of uncoupling DNA damage from replication stress in most eukaryotic models.

To investigate the temporal progression of genome instability in non-dividing cells, we used *Schizosaccharomyces pombe*, which enters a tightly regulated quiescent state upon nitrogen deprivation. We first examined cell viability over time in wild-type (WT) and selected DNA repair mutants aged up to 8 days in G0. While WT cells retained high viability, mutants lacking base excision repair (BER) enzymes (*ung1Δ, nth1Δ*), transcription-associated repair (*thp1Δ*), or combined deletions exhibited significantly accelerated viability loss, especially in the *nth1Δ ung1Δ thp1Δ* triple mutant (Figure 1A). This suggests that multiple repair systems act in parallel to preserve genome integrity and cellular fitness during quiescence.

**Figure 1.**
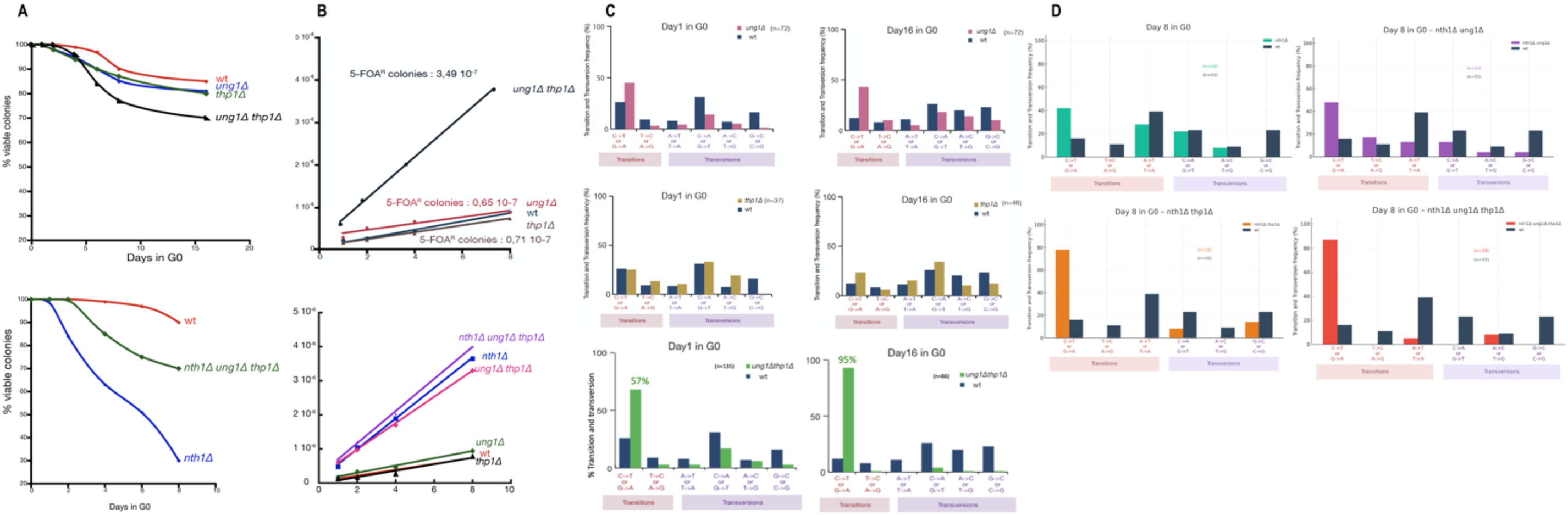
Viability loss, mutation accumulation, and distinct mutational spectra in quiescent S. pombe. **(A)** Cell viability of wild-type (WT) and repair-deficient mutants aged in G0 for up to 8 days. Single and combined mutants exhibit accelerated decline compared to WT. **(B)** Frequency of FOA-resistant (FQAAR) colonies increases progressively during quiescence, with higher accumulation in base excision repair-deficient *(ung1Δ, nth1Δ)* and transcription-associated repair mutants*(thp1Δ)*. **(C)** Mutation spectra of *ung1Δ, thp1Δ,* and the double mutant *(ung1Δ thp1Δ)* reveal enrichment in GC→AT transitions (uracil-driven) and clustered mutations in *thp1Δ,* consistent withtranscription-associated genome instability. **(D)** Mutation spectra of *nth1Δ, nth1Δ ung1Δ, nth1f1 thp1Δ,* and the *nth1Δ ung1Δ thp1Δ* triple mutant. The triple mutant displays the broadest spectrum, combining oxidative transversions withuracil- and transcription-linked signatures,indicative of synthetic genome instability in G0.

Next, we monitored the spontaneous accumulation of mutations by measuring the emergence of 5-fluoroorotic acid–resistant (FOAᴿ) colonies targeting the *ura4* and *ura5* loci. In WT cells, FOAᴿ mutants accumulated linearly with time, consistent with a basal mutagenesis rate in G0 (Figure 1B). However, the frequency of FOAᴿ colonies was markedly elevated in *ung1Δ, nth1Δ, thp1Δ*, and especially in the double and triple mutants, suggesting additive or synergistic effects of unrepaired base lesions and transcriptional stress.

To dissect the nature of accumulated mutations, we sequenced the *ura4* and *ura5* loci from FOAᴿ clones in each mutant background. In *ung1Δ*, the spectrum was dominated by GC→AT transitions, consistent with uracil misincorporation or cytosine deamination (Figure 1C). *thp1Δ* showed enrichment in clustered SNPs and complex base changes, likely linked to transcription-replication conflicts or persistent R-loops. In the *ung1Δ thp1Δ* double mutant, both signatures were present, indicating cooperation between uracil processing and R-loop resolution.

A more complex pattern emerged in *nth1Δ* and its derivatives (Figure 1D). While *nth1Δ* primarily showed oxidative base transversions (e.g., C→A, G→T), combining it with *ung1Δ* or *thp1Δ* led to broadened spectra. The *nth1Δ ung1Δ thp1Δ*triple mutant accumulated both oxidative and uracil-associated mutations, as well as compound indels and clustered substitutions - hallmarks of synthetic genome instability. This combinatorial mutational landscape reveals that multiple endogenous lesion types accumulate in G0 and that their repair is not functionally redundant.

Together, these results indicate that quiescent cells experience progressive viability loss and accumulate mutations in a lesion- and pathway-specific manner. The mutation spectrum is strongly shaped by the activity of DNA glycosylases, transcription-associated processing factors, and their combinatorial interactions. These findings establish the foundation for our further dissection of how repair pathways cooperatively maintain genome stability during cellular dormancy.

### 2. Uracil is a major endogenous source of mutagenesis in quiescent BER-deficient cells

The mutational spectra identified in the previous section revealed that GC→AT transitions represent a prominent signature in *ung1Δ* and combined BER-deficient mutants aged in quiescence (Figure 1C–D). These mutations are strongly suggestive of uracil accumulation in DNA, either through spontaneous cytosine deamination or dUTP misincorporation during DNA repair synthesis. Under physiological conditions, such lesions are repaired through the base excision repair (BER) pathway, initiated by uracil-DNA glycosylases such as Ung1, followed by AP site processing via Nth1 [5,6,16,17].

To directly test whether uracil drives G0-associated mutagenesis, we quantified mutation rates and lesion signatures in *nth1Δ, ung1Δ*, and *nth1Δ ung1Δ* double mutants after 8 days in G0. These strains showed a sharp increase in FOAᴿ colony frequency compared to WT, and their mutational spectra were enriched in C: G > T:A transitions, a hallmark of uracil-driven mutagenesis (see Figure 1C–D). These patterns confirm that uracil-related damage becomes a major driver of genome instability when the BER pathway is impaired during quiescence.

To determine whether uracil physically accumulates in the genome during quiescence, we employed three complementary and orthogonal approaches. First, an alkaline cleavage assay using recombinant *E. coli* Ung1 revealed pronounced fragmentation of genomic DNA extracted from the *nth1Δ ung1Δ thp1Δ* triple mutant, consistent with enzymatic conversion of uracil into single-strand breaks (Figure 2A) [44]. Second, a dot blot assay using a uracil-specific monoclonal antibody detected elevated uracil signals in DNA from BER-deficient strains, with minimal signal observed in wild-type cells (Figure 2B). Finally, LC–MS/MS analysis of genomic DNA confirmed the chemical presence of uracil in quiescent cells, with significantly higher levels detected in *nth1Δ ung1Δ* mutants compared to WT (Figure 2C). Together, these results provide direct molecular evidence that uracil lesions accumulate in the genome during quiescence, particularly in the absence of a functional base excision repair pathway.

**Figure 2.**
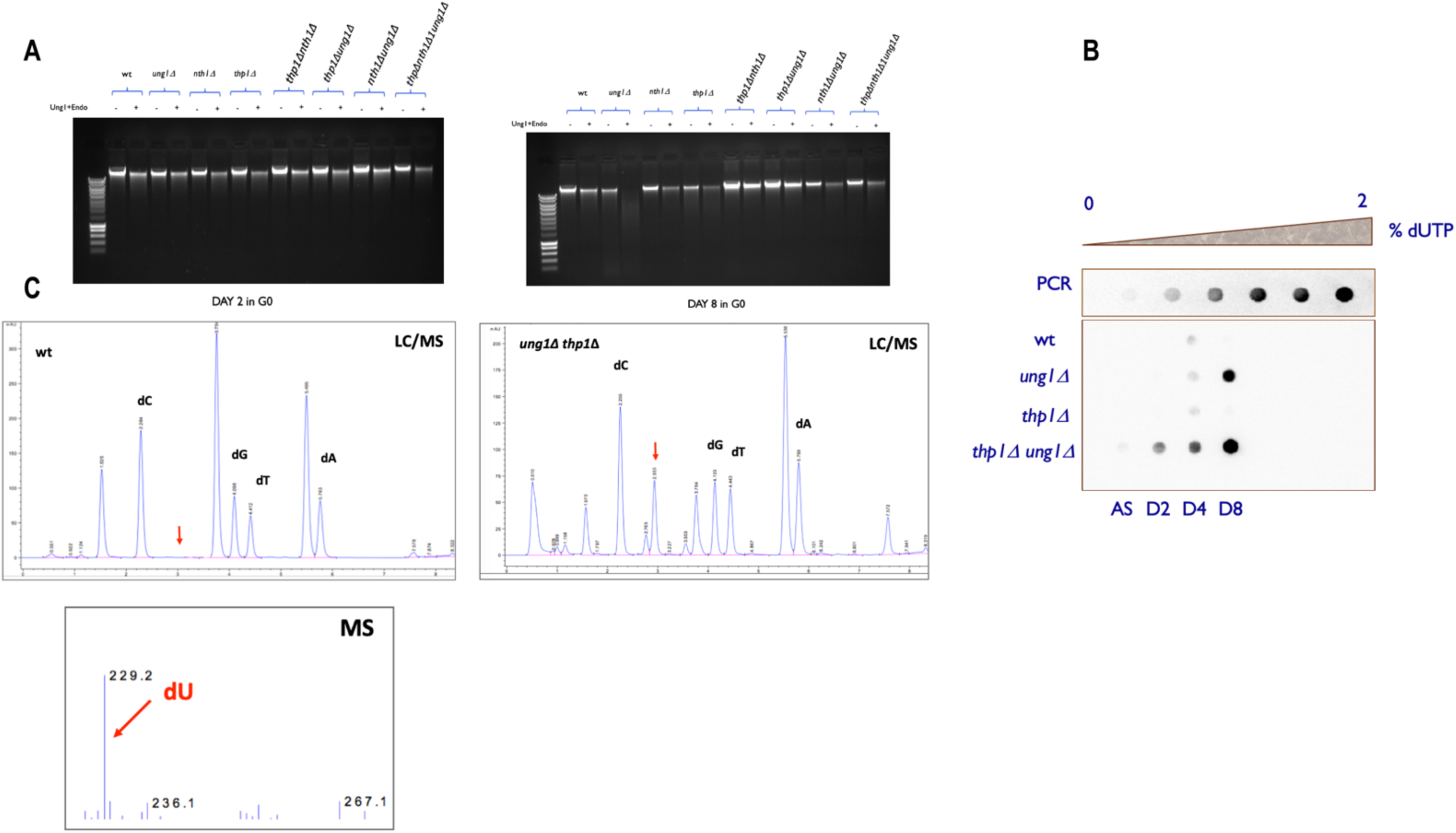
Uracil accumulation in quiescent BER-deficient cells. **(A)** Alkaline cleavage assay of genomic DNA extracted from wild-type (WT) and *nth1Δ ung1Δ thp1Δ* triple mutant cells aged in quiescence (day 8), incubated with recombinant *E. coli* Ung1. The increased DNA fragmentation observed in the mutant reflects the accumulation of unrepaired uracil residues. **(B)** Dot blot analysis using a uracil-specific monoclonal antibody further confirms the higher uracil content in the triple mutant compared to WT. **(C)** Detection of free uracil by liquid chromatography-mass spectrometry (LC-MS/MS) confirms the biochemical accumulation of uracil in genomic DNA from the *ung1Δ thp1Δ* mutant, further validating the role of BER components in limiting uracil persistence during quiescence.

To explore whether uracil accumulation is exacerbated by metabolic imbalance, we treated quiescent WT cells with methotrexate (MTX) to artificially perturb nucleotide pools. MTX inhibits dihydrofolate reductase, reducing dTTP synthesis and increasing the dUTP/dTTP ratio, which promotes uracil misincorporation. Consistent with this, MTX-treated cells showed increased DNA fragmentation following Ung1 treatment (Figure 4A) and a rise in FOAᴿ mutagenesis (Figure 4B), even in the WT background. These results suggest that nucleotide pool imbalance, a hallmark of nutrient deprivation, may further contribute to uracil-driven damage in G0 [56,37].

Together, these findings demonstrate that uracil is a major endogenous DNA lesion in non-dividing cells, particularly in the context of impaired BER. The data also suggest that dUTP accumulation and cytosine deamination are major sources of genomic uracil during quiescence, and that metabolic stress exacerbates this process, ultimately leading to mutagenesis and genome instability.

### 3. Overlapping functions of Hnt3 and Rnh201 in suppressing quiescent genome instability

To further dissect the landscape of genome surveillance in G0, we investigated whether Hnt3 and Rnh201-enzymes involved in repair of 5′-end blocking lesions and R-loop resolution, respectively-act in redundant or complementary pathways during quiescence.

As shown in Figure 3A, both *hnt3Δ* and *rnh201Δ* single mutants exhibited modest but significant increases in FOA-resistant mutant frequency after 8 days in G0, consistent with accumulation of unrepaired spontaneous lesions. However, the *hnt3Δ rnh201Δ* double mutant showed a synergistic elevation in mutagenesis, exceeding the additive effect of single mutants. This suggests that these enzymes operate in partially overlapping but distinct genome protection pathways.

**Figure 3.**
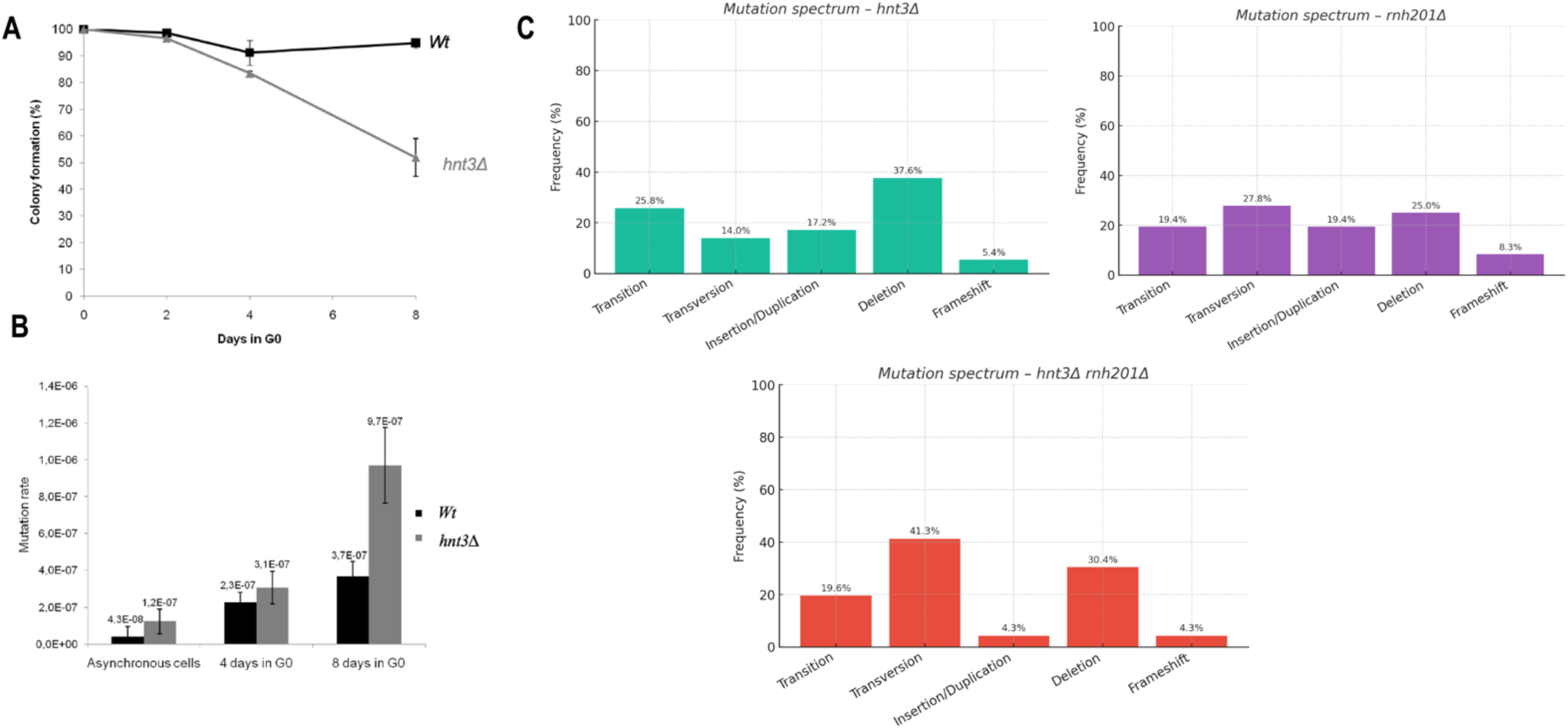
Hnt3 and Rnh201 act in distinct but partially overlapping pathways to prevent quiescent genome instability. **(A)** Frequency of FOA-resistant mutants in *hnt3Δ, rnh201Δ,* and *hnt3Δ rnh201Δ* strains aged for 8 days in G0. A synergistic increase in mutagenesis is observed in the double mutant, indicating cooperation between the two pathways. **(B)** Quantification of colony morphologies upon quiescence exit. The *hnt3Δ rnh201Δ* mutant shows a reduction in viable colonies and an increase in microcolony/G0-like structures, suggesting post-quiescence growth defects. **(C)** Mutation spectrum in ura 4 and ura 5 loci from FOA-resistant clones. The double mutant accumulates more indels and complex mutations than either single mutant.

**Figure 4.**
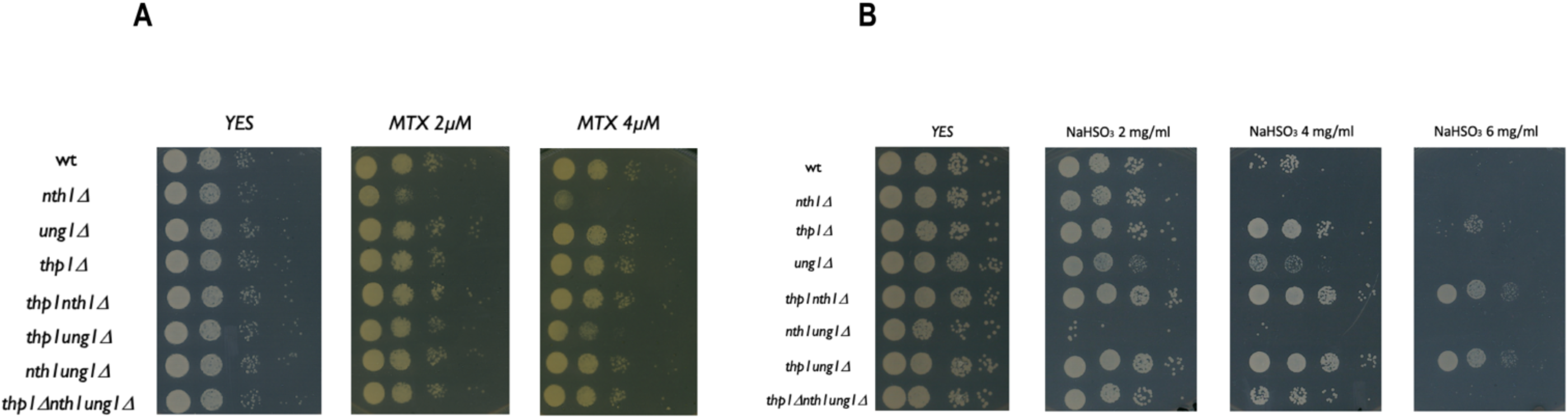
Nucleotide pool imbalance and oxidative base damage synergize to reveal BER dependency in quiescent cells. **(A)** Spot dilution assays of wild-type and BER-deficient mutants *(nth1Δ, ung1Δ, thp1Δ,* and combinations) aged for 8 days in quiescence and plated on YES medium supplemented with methotrexate (MTX; 2µM and 4µM), which inhibits thymidylate synthesis and induces dUTP/dTTP imbalance. The triple mutant *thp1Δ nth1Δ ung1Δ* exhibits a pronounced growth defect under MTX, indicating hypersensitivity to uracil misincorporation and defective base excision repair. **(B)** The same set of strains was tested for sensitivity to sodium bisulfite (NaHS0_3_; 2-6mg/ml), a chemical that induces cytosine deamination and abasic site formation. Progressive loss of viability is observed with increasing NaHS0_3_ concentrations, particularly in *nth1Δ, ung1Δ,* and the triple mutant background, consistent with impaired processing of abasic and oxidized lesions.

Analysis of post-quiescence colony formation (Figure 3B) revealed that the double mutant was also severely impaired in recovery, with a higher proportion of microcolonies and G0-like phenotypes, indicating incomplete cell cycle re-entry and likely persistent damage.

Moreover, the mutation spectrum in *hnt3Δ rnh201Δ* clones (Figure 3C) was skewed toward small deletions, microduplications, and complex indels, implying unresolved DNA ends or persistent transcription-associated damage. These mutation patterns are consistent with previous observations in *S. cerevisiae* and mammalian systems, where loss of APTX (the human homolog of Hnt3) or RNase H2 leads to genome instability, neurological disorders, and inflammatory phenotypes due to endogenous DNA damage and RNA: DNA hybrid accumulation [39,40,19,33,47].

Together, these findings reinforce the idea that Hnt3 and Rnh201 function in parallel to suppress spontaneous DNA damage during quiescence. Their synthetic phenotypes suggest that quiescent genome maintenance requires coordinated resolution of abortive repair intermediates and R-loop-mediated stress to prevent mutagenesis and ensure proper re-entry into the cell cycle.

### 4. mre11Δ cells exhibit reduced quiescent viability and accumulate multiple mutations per locus

The Mre11-Rad50-Nbs1 (MRN) complex plays a key role in the repair of DNA double-strand breaks (DSBs), replication fork stabilization, and checkpoint activation. However, its contribution to genome maintenance in non-dividing, quiescent cells remain largely unexplored. To address this, we assessed the behavior of *mre11Δ* mutants during prolonged G0 arrest.

First, we measured cell viability after 8 days in G0. While WT cells maintained high viability (∼90%), *mre11Δ* cells showed a significant reduction in colony formation (∼55%), indicative of cumulative damage during quiescence that impairs regrowth capacity (Figure 5A). Consistently, we observed an increased frequency of FOA-resistant mutants in *mre11Δ* compared to WT (Figure 5B), suggesting enhanced mutagenesis in the absence of Mre11 function. To characterize the nature of these mutations, we sequenced the *ura4* and *ura5* loci in independent FOAᴿ *mre11Δ* clones. Mutations were distributed throughout both genes without obvious hotspots (Figure 5C). Strikingly, several individual clones harbored multiple independent mutations within a single gene (Figure 5D), an observation rarely seen in WT or other mutants. These complex mutation patterns likely reflect the inability of *mre11Δ* cells to properly resolve spontaneous DNA breaks or replication-associated intermediates during quiescence. A detailed analysis of mutation types revealed a higher proportion of small insertions/deletions (1–5 bp) and complex indels in *mre11Δ*, as shown in the aggregated mutational spectrum (Figure 5E). Furthermore, time-course experiments showed a progressive shift from base substitutions at day 1 to an enrichment of indels and complex events at later time points (day 8 and day 15), suggesting a cumulative failure in genome maintenance (Figure 5F).

**Figure 5.**
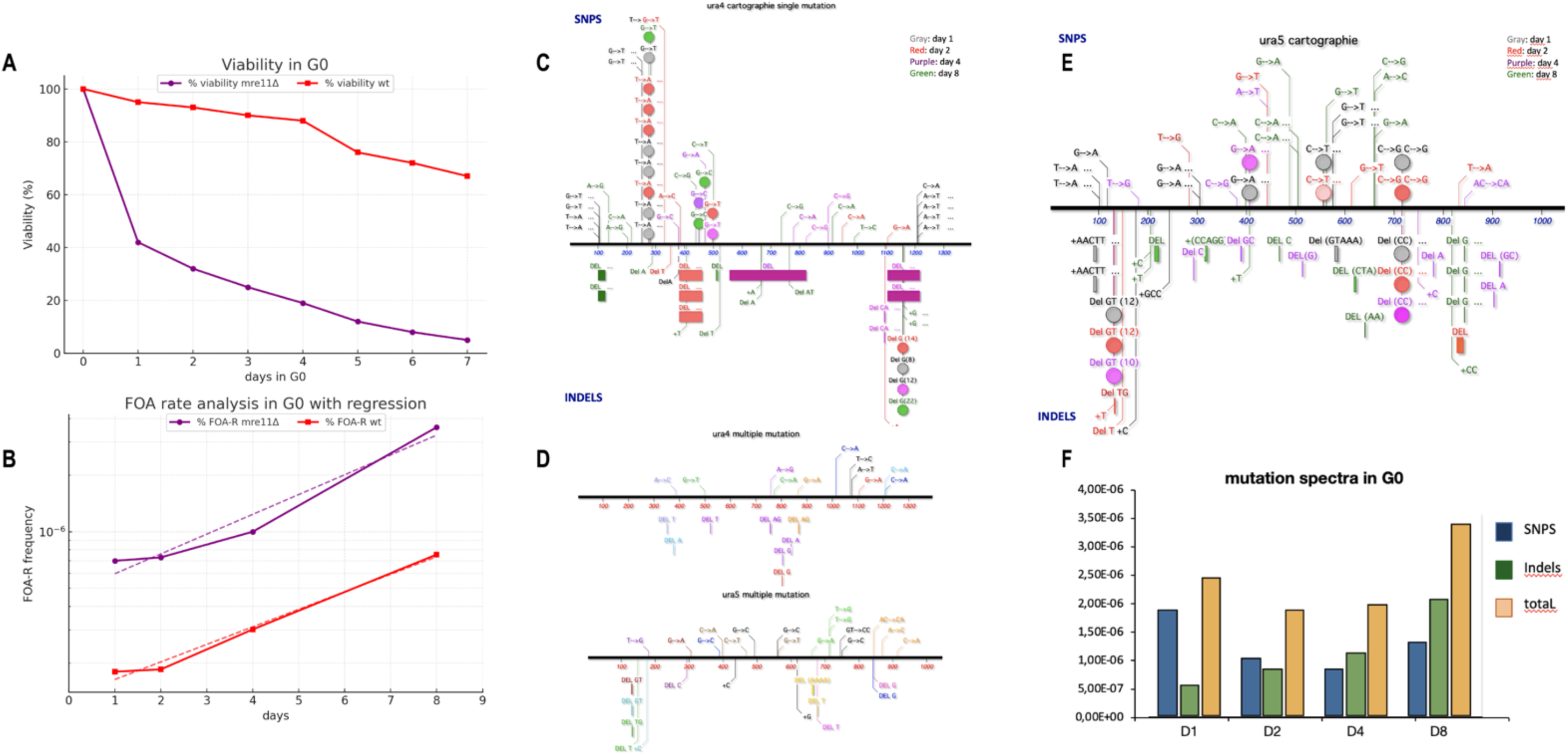
Quiescent mre11Δ cells exhibit increased mutagenesis, decreased viability, and complex mutation patterns. **(A)** Quantification of viability after 8 days in G0 for mre11Δ and WT cells. mre11Δ cells exhibit significantly reduced colony-forming capacity upon exit from quiescence. **(B)** FOA-resistant mutant frequency in mre11Δ compared to WT, revealing elevated mutation accumulation during quiescence. **(C)** Mutation mapping in the ura4 and ura5 loci of FOA^R mre11Δ clones, indicating random distribution of mutations across both genes. **(D)** High-resolution mapping of multiple mutations per clone in ura4 and ura5 in mre11Δ, showing frequent accumulation of ≥2 mutations per gene in single cells. **(E)** Pie chart illustrating the distribution of mutation types (SNPs, small indels, large deletions, complex events) in mre11Δ across multiple time points. **(F)** Mutation spectrum in mre11Δ over increasing durations in G0 (days 1, 2, 4, 8), showing a time-dependent shift from base substitutions to complex structural variants.

Together, these findings establish that Mre11 is essential to suppress spontaneous DNA damage and maintain genome stability in quiescent cells. The increased mutation load, combined with reduced viability and accumulation of multiple hits per locus, point to a critical role for the MRN complex in the long-term survival and genetic integrity of non-dividing cells [53].

### 5. Synthetic interactions reveal pathway crosstalk in post-quiescence recovery

The accumulation of mutations and viability loss in *mre11Δ* prompted us to explore how other repair pathways interact functionally with Mre11 in quiescent cells. To dissect potential pathway redundancy or compensation, we generated double mutants combining *mre11Δ* with deletions in genes involved in single-strand break (SSB) repair (*tdp1Δ*), homologous recombination (*rhp52Δ*), and nucleotide excision repair (*rad13Δ*). We then monitored their post-quiescence recovery by analyzing colony morphology and viability on solid medium after 8 days in G0.

Using a single-cell dissection approach, we followed the fate of individual cells plated on YES medium upon exit from G0. This technique revealed significant heterogeneity in colony-forming capacity among mutants (Supplementary Figure S1). While WT and *mre11Δ* cells were still able to form full colonies in most cases, the double mutants *mre11Δ tdp1Δ*and *mre11Δ rhp52Δ* exhibited a marked shift toward microcolony formation, indicating incomplete or delayed recovery.

To quantify these phenotypes, we analyzed the percentage of viable colonies per genotype and scored colony size distributions. The results are summarized in Supplementary Figure S2, which highlights the synthetic defects in double mutants. Notably, only ∼40% of *mre11Δ rhp52Δ* cells formed viable colonies, compared to ∼90% in WT and ∼55% in *mre11Δ* alone. The synthetic interaction between *mre11Δ* and *rhp52Δ* suggests that homologous recombination compensates for Mre11 loss in repairing spontaneous DSBs during quiescence. A similar, though less severe, trend was observed for *mre11Δ tdp1Δ*, implying a partial role for SSB repair in buffering Mre11 deficiency [41,48].

In contrast, the *mre11Δ rad13Δ* combination did not show a significant decrease in viability, suggesting that nucleotide excision repair plays a lesser role in the quiescent context. These observations underscore a hierarchy of repair pathway importance during G0, with Mre11 and Rhp52 playing dominant roles in survival and genome maintenance.

Together, these results demonstrate that loss of Mre11 triggers a reliance on backup pathways, particularly homologous recombination, to ensure successful quiescence exit. The failure of these compensatory mechanisms in double mutants results in synthetic growth defects and increased mutagenic load.

### 6. Single-cell resolution reveals synthetic defects in DNA repair double mutants

To better resolve the cellular consequences of endogenous DNA damage during quiescence, we employed a single-cell dissection assay, allowing us to monitor individual cells aged in G0 as they attempt to re-enter the cell cycle upon re-feeding.

As shown in Supplementary Figure S1, WT cells retained robust proliferation capacity and formed mature colonies. In contrast, several double mutants - notably *hnt3Δ rhp52Δ, tdp1Δ rhp52Δ,* and *sen1Δ rad2Δ* - showed defective recovery, forming only microcolonies or remaining in a G0-like arrested state. These phenotypes reflect persistent DNA lesions incompatible with full proliferation.

We quantified the outcomes by classifying colonies into three categories: mature (viable), microcolonies (≤8 cells), and arrested (no division). The distribution of these classes (Supplementary Figure S2) confirmed that *hnt3Δ rhp52Δ* and *tdp1Δ rhp52Δ* have severely compromised quiescence exit, consistent with synthetic defects.

These data strongly suggest that genome maintenance in quiescence depends on coordinated repair pathways, and that the combined inactivation of DNA end-processing factors (e.g., Tdp1, Hnt3) with recombination (e.g., Rhp52) or NER (e.g., Rad2) produces unrecoverable lesions [39,41,47,64].

## Discussion

Maintaining genome integrity in non-dividing cells is a fundamental challenge for long-term cellular survival and organismal health. Using *Schizosaccharomyces pombe* as a model, we investigated how the genome is reshaped and preserved during prolonged quiescence induced by nitrogen starvation. Our study demonstrates that quiescent cells, although non-replicating, accumulate spontaneous DNA lesions that lead to mutagenesis, and that this process is tightly regulated by a coordinated network of repair pathways.

We first established that quiescent wild-type cells gradually accumulate mutations over time, with a clear transition from single-nucleotide substitutions to complex structural variants (Figure 1). This pattern reflects a time-dependent accumulation of spontaneous damage, consistent with previous reports in *S. cerevisiae* and mammalian systems suggesting ongoing genome reshaping during G0 arrest [7,52]. Despite this increase in mutation load, WT cells maintained high viability, suggesting the presence of efficient surveillance and repair systems even in the absence of DNA replication.

A key finding of this study is the identification of uracil as a major endogenous mutagen in G0. Through a combination of alkaline cleavage assays, antibody-based detection, and direct LC–MS/MS quantification, we confirmed that uracil accumulates in the genome of BER-deficient cells aged in G0 (Figure 2). This lesion likely arises from both cytosine deamination and dUTP misincorporation under conditions of nucleotide imbalance - a scenario exacerbated by starvation. The resulting C: G > T: A transitions and small deletions in *nth1Δ ung1Δ* mutants mirror classical uracil-driven mutagenesis signatures [5,16,17,36]. These results place base excision repair (BER), particularly the Nth1–Ung1 axis, at the forefront of genome protection during quiescence.

Moreover, our data show that perturbing dNTP homeostasis with methotrexate further enhances uracil accumulation and FOA resistance (Figure 4), supporting the notion that nucleotide pool imbalance is a driver of G0 genome instability [37,56]. This suggests potential metabolic vulnerabilities in quiescent cells that could be exploited in therapeutic contexts, such as cancer cell dormancy.

We also uncovered a prominent role for the DNA end-processing factor Mre11 in G0 genome maintenance. *mre11Δ* cells displayed elevated mutagenesis, reduced viability, and the frequent accumulation of multiple mutations per clone (Figure 5). The shift toward complex indels and the temporal diversification of mutational spectra support a model in which Mre11 acts as a key suppressor of replication-independent DNA damage. This expands its classical role in DSB repair to include broader genome surveillance functions in non-cycling cells [3,53].

Importantly, genetic interaction analyses revealed that Mre11 collaborates with both homologous recombination and SSB repair pathways to ensure survival and genome integrity during G0. Double mutants combining *mre11Δ* with *rhp52Δ* or *tdp1Δ* showed synthetic defects in post-quiescence regrowth, indicating pathway interdependence (Supplementary Figures S1–S2). These findings mirror observations in *S. cerevisiae* and metazoan cells, where dormant cells often rely on HR-mediated repair to resolve transcription-associated damage or persistent oxidative lesions [3,48].

In parallel, our study also implicates transcription-associated genome instability as a significant source of quiescent mutagenesis. The RNA/DNA helicase Sen1, previously associated with R-loop resolution, was shown to play a key role in protecting the genome during G0. The *sen1Δ* mutant displayed a marked increase in mutation frequency and a shift in mutational spectrum, consistent with the accumulation of transcription-associated lesions. Notably, genetic interaction studies revealed synthetic defects in *sen1Δ rad2Δ* and *sen1Δ rhp52Δ* mutants, suggesting that unresolved R-loops may be converted into DSBs that require homologous recombination or nucleotide excision repair for resolution. These findings are consistent with studies in human and yeast cells linking SEN1/Senataxin deficiency to increased transcriptional stress and genome instability [20,33,21,25].

In addition to DSB and BER pathways, we identified a cooperative role between the R-loop processing factor Rnh201 and the DNA end-cleaning enzyme Hnt3. The *hnt3Δ rnh201Δ* double mutant exhibited a synthetic increase in mutation load and post-quiescence growth defects, accompanied by a distinct mutational profile rich in indels and complex rearrangements. These findings are reminiscent of the genomic instability observed in RNase H2- or aprataxin-deficient mammalian cells and underscore the importance of RNA: DNA hybrid resolution and DNA end-processing in the maintenance of genome integrity in non-replicating states [39,19,47,64].

Altogether, our results highlight the unique landscape of genome maintenance in quiescent cells, involving uracil-driven mutagenesis, transcription-associated stress, and reliance on coordinated repair networks including BER, DSB repair, and SSB processing. The synthetic interactions uncovered in double mutants emphasize the importance of pathway redundancy in protecting genome integrity in non-dividing states.

Our single-cell assay revealed phenotypes that bulk FOA mutagenesis failed to capture. While FOA resistance quantifies fixed mutations, colony dissection provides insight into transient or sublethal DNA damage. The synthetic defects in *hnt3Δ rhp52Δ* and *tdp1Δ rhp52Δ* underscore the importance of cooperative repair mechanisms in G0. Persistent DNA ends or unresolved R-loops likely interfere with replication restart, leading to incomplete cell cycle re-entry. These findings echo studies in post-mitotic mammalian cells, where similar defects underlie neurodegeneration and immune failure [31,60,50].

Understanding how quiescent cells manage genome stability has broad implications, from aging and stem cell biology to cancer therapy, where dormant cells often evade genotoxic stress. Our findings offer a mechanistic framework for future studies into quiescent genome dynamics and the vulnerabilities of non-replicating cells [51,62].

In conclusion, our study reveals that genome maintenance in quiescent *Schizosaccharomyces pombe* cells is far from passive and instead relies on a dynamic interplay of DNA repair pathways that differ markedly from those active in proliferating cells. By longitudinally tracking mutation accumulation during nitrogen starvation-induced G0, we demonstrate that quiescent cells undergo a progressive and significant rise in mutational burden, including complex indels and structural variants.

We identify uracil misincorporation and deamination as major endogenous sources of G0 mutagenesis, particularly exacerbated in base excision repair (BER)-deficient backgrounds. Moreover, the data underscore the critical role of Mre11, Tdp1, Hnt3, and Sen1 in restraining DNA damage in non-dividing cells. Their genetic interactions with homologous recombination or nucleotide excision repair components reveal parallel and compensatory repair routes required for successful re-entry into the cell cycle.

Importantly, high-resolution single-cell analyses uncovered latent repair defects not detectable in bulk assays. Strains such as *hnt3Δ rhp52Δ* and *tdp1Δ rhp52Δ* displayed profound recovery failures, illustrating how genome instability in quiescence can impact long-term viability and proliferation potential.

Collectively, our work defines a new framework for understanding genome integrity in non-cycling cells. These findings may have broad implications for quiescent stem cells, immune memory, and aging, where low-level DNA damage accumulates silently over time but becomes biologically consequential upon activation or replication re-entry.

## Data Origin, Figures, and Materials Availability

The data presented in this manuscript were generated between 2009 and 2015 as part of an independent research initiative that I conceived and developed during my doctoral and postdoctoral training. Although these results were never formally published, they remain highly relevant to the field of genome stability in non-dividing cells. By depositing this work as a preprint on bioRxiv, I aim to ensure the permanent accessibility of these findings to the scientific community and to establish an official, traceable record of authorship.

All data, figures, and interpretations presented in this preprint are the intellectual property of the author. This work is released under the Creative Commons Attribution (CC BY 4.0) license selected during submission. Any reproduction, reuse, or derivative work - including figures, datasets, or textual excerpts - must properly cite this preprint using its DOI and full bibliographic reference. Unauthorized use without attribution constitutes a violation of copyright and of the terms of this license.

## Acknowledgments

This study includes experimental work initiated and developed by the author between 2009 and 2015 during doctoral and postdoctoral training in the laboratory of Benoit Arcangioli at the Pasteur Institute. The project was independently conceived and carried out by the author, who remains solely responsible for the data, analyses, and interpretations presented in this manuscript.

Portions of the work were conducted using shared laboratory infrastructure, with technical access and materials provided by the Arcangioli lab. The author is grateful for this support and acknowledges the helpful contributions of former lab members during the early development of the project.

The author also wishes to sincerely thank Dr. Sylvie Pochet and Valérie Huteau from the Chemistry Unit at the Pasteur Institute for their valuable assistance with LC/MS and MS analyses, which greatly enhanced the biochemical validation of uracil accumulation.

## Supplementary Information

**Supplementary Figure S1.**
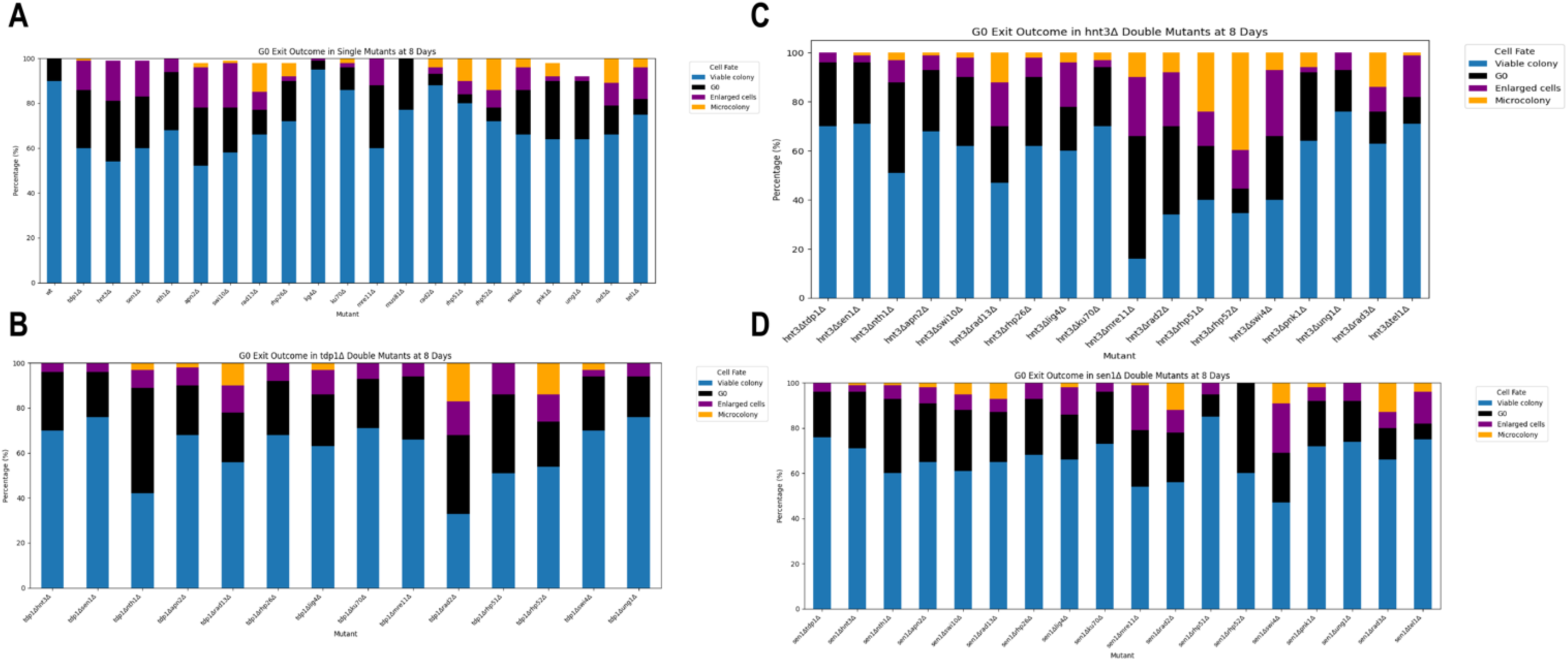
Exit from G0 in single and double mutants involving *hnt3Δ, sen1Δ,* and *tdp1Δ.* Stacked bar charts representing the cellular outcomes following G0 exit at day 8 for the indicated mutants. Each bar corresponds to one strain and displays the proportion of cells classified as viable colonies (blue), microcolonies (orange), enlarged unbudded cells (purple), or arrested in G0 (black). Strains shown include: **(A)** single mutants, **(B)** *tdp1Δ* double mutants, **(C)** *hnt3Δ* double mutants, and **(D)** *sen1Δ* double mutants. Data were obtained from single-cell analyses using micromanipulation of individual G0 cells on YES plates and phenotypic classification after 5 days at 32°C. Results highlight genetic interactions affecting G0 exit efficiency and cellular fate.

**Supplementary Figure S2.**
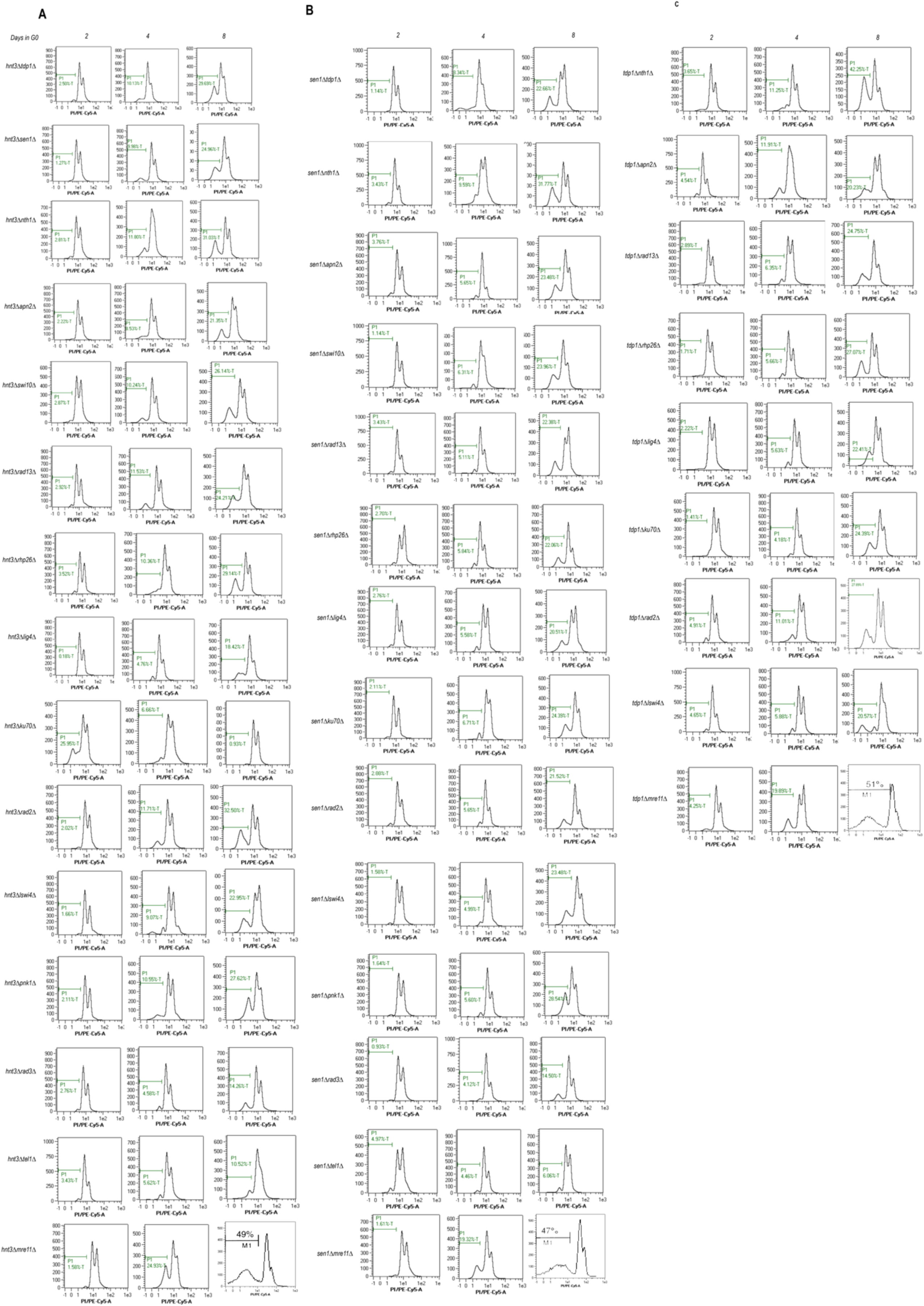
Flow cytometry analysis of G0-phase cell cycle profiles in double DNA repair mutants. **(A)** hnt3Δ-based double mutants (APT series): Flow cytometry DNA content profiles (Pl staining) of hnt3Δ combined with repair mutants (apn2Δ, nth1Δ, rhp26Δ, swi10Δ, etc.) after 2, 4, and 8 days in quiescence. Some combinations exhibit moderate G2/M accumulation or DNA fragmentation. **(B)** sen1Δ-based double mutants (SEN series): Profiles of sen1Δ combined with mre11Δ, rad13Δ, ku70Δ, rhp26Δ, etc. show marked sub-G1 peaks and delayed exit from G0, indicating persistent DNA damage and checkpoint activation. **(C)** Tdp1Δ-based double mutants (TOP series): Flow cytometry profiles of tdp1Δ combined with swi10Δ, rad13Δ, lig4Δ, rad2Δ, and others. Many shows abnormal DNA content distribution, sub-G1 accumulation, or G2/M delay, reflecting defective repair coordination in quiescence. Note: Cells were stained with propidium iodide after ethanol fixation and analyzed on a BO Accuri C6 cytometer. Histograms represent DNA content distribution at each time point.

